# Interspecies signaling generates exploratory motility in *Pseudomonas aeruginosa*

**DOI:** 10.1101/607523

**Authors:** Dominique H. Limoli, Niles P. Donegan, Elizabeth A. Warren, Ambrose L. Cheung, George A. O’Toole

## Abstract

Microbes often live in multispecies communities where interactions among community members impact both the individual constituents and the surrounding environment. Here, we developed a system to visualize interspecies behaviors at initial encounters. By imaging two prevalent pathogens known to be coisolated from chronic illnesses, *Pseudomonas aeruginosa* and *Staphylococcus aureus*, we observed *P. aeruginosa* can modify surface motility in response to secreted factors from *S. aureus*. Upon sensing *S. aureus, P. aeruginosa* transitioned from collective to single-cell motility with an associated increase in speed and directedness – a behavior we refer to as ‘exploratory motility’. Through modulation of cAMP, explorer cells moved preferentially towards *S. aureus* and invaded *S. aureus* colonies through the action of the type IV pili. These studies reveal previously undescribed motility behaviors and lend insight into how *P. aeruginosa* senses and responds to other species. Identifying strategies to harness these interactions may open avenues for new antimicrobial strategies.

## Introduction

It has been clear since the 1600’s that many microbial infections do not occur with a single species, but we have only recently begun to understand the profound impacts microbial species have on each other and patients (Nguyen and Oglesby-Sherrouse 2016)}. Studies of dental biofilms, intestinal communities, chronic wounds, and respiratory infections in patients with cystic fibrosis (CF) demonstrate that community interactions influence microbial survival and disease progression (Limoli and Hoffman 2019; Gabrilska and Rumbaugh 2015). For example, we and others find an association between coisolation of *Pseudomonas aeruginosa* and *Staphylococcus aureus* from the CF airway or chronic wounds and poor patient outcomes, including decreased lung function and shortened life-spans (Limoli et al. 2016; Maliniak, Stecenko, and McCarty 2016; Hubert et al. 2013). Laboratory studies also reveal interactions between these two pathogens can alter virulence factor production by one or both species, potentially influencing pathogenesis, persistence, and/or antibiotic susceptibility (Hotterbeekx et al. 2017). For example, in a model of coinfection on CF-derived bronchial epithelial cells, we observed *P. aeruginosa* and *S. aureus* form mixed microcolonies, which promotes the survival of *S. aureus* in the presence of vancomycin (Orazi and O’Toole 2017). One strategy to improve outcomes for these coinfected patients is to block harmful interspecies interactions before they begin.

Here, we designed a system to visualize early interactions between *P. aeruginosa* and *S. aureus*, and follow single-cell behaviors over time with live imaging. We show that *P. aeruginosa* can sense *S. aureus* secreted products from a distance and in turn, dramatically alter the motility behaviors of this Gram-negative bacterium. In response to *S. aureus*, individual *P. aeruginosa* cells transition from collective to single cell movement, allowing exploration of the surrounding environment and directional movement towards *S. aureus.* We find that such ‘exploratory motility’ is driven primarily by the *P. aeruginosa* type IV pili and modulation of the intracellular second messenger, cAMP. Thus, we provide a new means to study polymicrobial interactions at the single-cell level and reveal that *P. aeruginosa* can sense the presence of other microbial species and dramatically, yet specifically, modify its behavior in response to such interspecies signals.

## Results

### *S. aureus* promotes exploratory motility in *P. aeruginosa*

To understand early microbial interactions, we established an *in vitro* coculture system to monitor *P. aeruginosa* and *S. aureus* at first encounters. Bacteria were inoculated at low cell densities between a coverslip and an agarose pad in minimal medium, supplemented with glucose and tryptone, and imaged with phase contrast time-lapse microscopy every 15 minutes for 8 hours. Alone, *P. aeruginosa* cells replicate and expand outward as raft-like groups, as previously described for *P. aeruginosa* surface-based motility (Anyan et al. 2014; Burrows 2012) (**Movie 1**; **Figure 1**, still montage, top row). In comparison, coincubation with *S. aureus* resulted in dramatically altered behavior (**Movie 2, Figure 1**, still montage, bottom row). After two to three rounds of cell division, instead of remaining as a group, individual *P. aeruginosa* cells began to move as single cells, suggesting that *P. aeruginosa* responds to the presence of *S. aureus* by altering motility behaviors. *P. aeruginosa* significantly inhibited *S. aureus* growth, as previously reported (see **Movie 3** for representative time-lapse movie of *S. aureus* alone).

**Figure 1.**
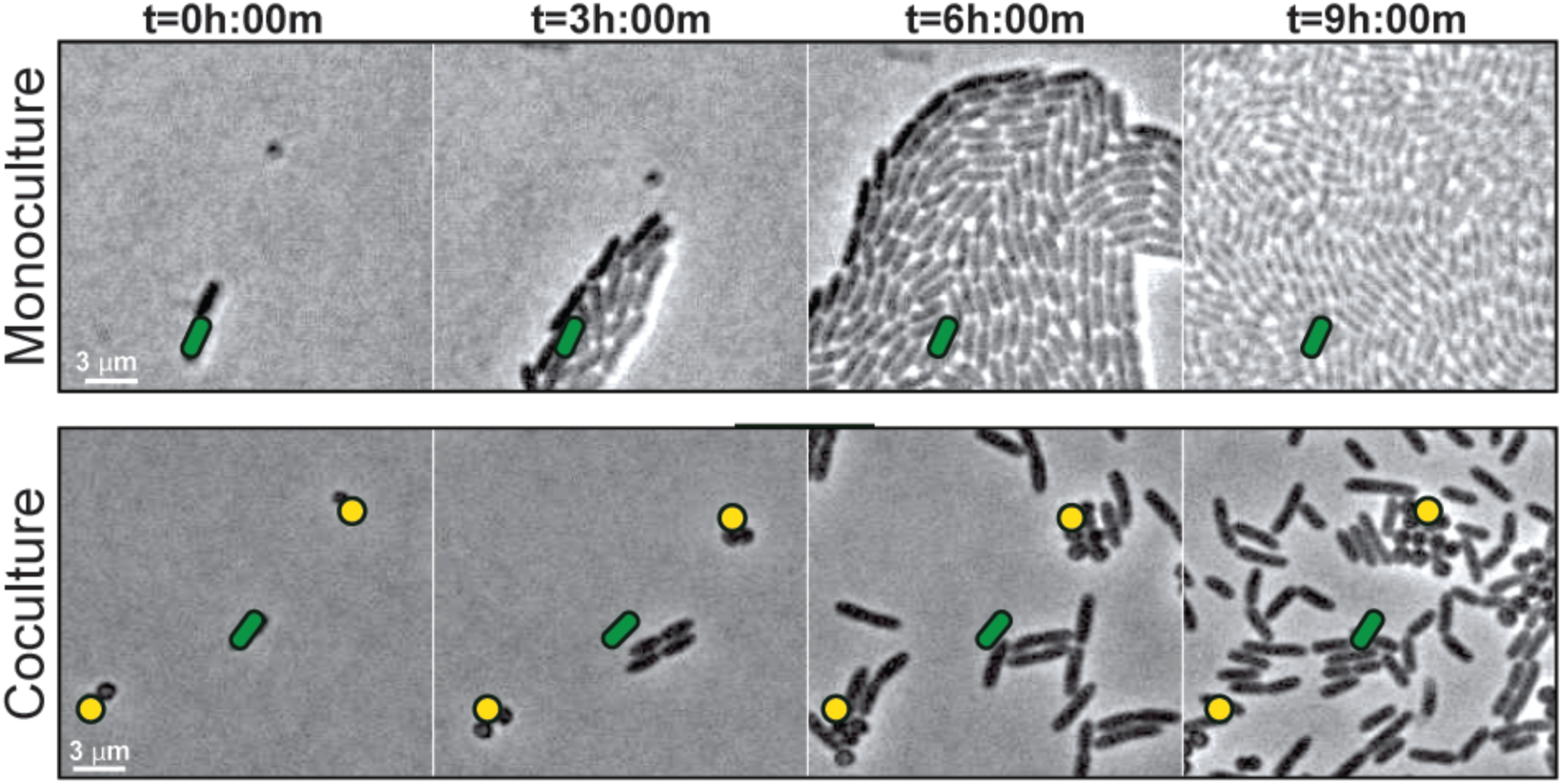
*S. aureus* increases *P. aeruginosa* motility. Live imaging of polymicrobial interactions. *P. aeruginosa* (rod-shaped) was inoculated between a coverslip and an agarose pad, either in monoculture (top) or in coculture with equal numbers of *S. aureus* (cocci-shaped, bottom). Images were acquired every 15m for 9h. Representative snap-shots of **Movies 1** (top) and **2** (bottom) are shown. Founding cells identified in the first frame are indicated with green rods (*P. aeruginosa*) or yellow circles (*S. aureus*). The location of the founding cell is indicated in each subsequent frame for positional reference.

To visualize *P. aeruginosa* motility in the presence of *S. aureus* in more detail, the inoculating cell density was reduced (2-3 cells of each species per field of view), and images were taken at 5s intervals for 8h. **Movie 4** shows images taken during hours 4 – 6 of coculture, when *P. aeruginosa* initiates single-cell movement under these conditions (**Figure 2A**, still montage). We observed a number of surprising behaviors by *P. aeruginosa* in the presence of *S. aureus. P. aeruginosa* cells initially replicated and remained in a raft (t=4h:28m), as we and others have observed for *P. aeruginosa* in monoculture, but as the community approached *S. aureus*, individual cells: (1) exit the raft (t=4h:34m), (2) move with increased speed, and (3) move towards and surround *S. aureus*, “exploring” the surface of the colony until (**Figure 2A-B**), (4) *P. aeruginosa* cells enter the *S. aureus* colony (**Figure 2C**, t=6h,), and finally, (5) by 8h, *P. aeruginosa* completely dismantles the *S. aureus* community (**Figure 2D**, t=8h). *P. aeruginosa* was also observed to adopt a swift motion, beginning between 4.5 - 6h, moving in and out of the plane of focus during imaging (**Movie 5** and **Figure 2D**, red arrows).

**Figure 2.**
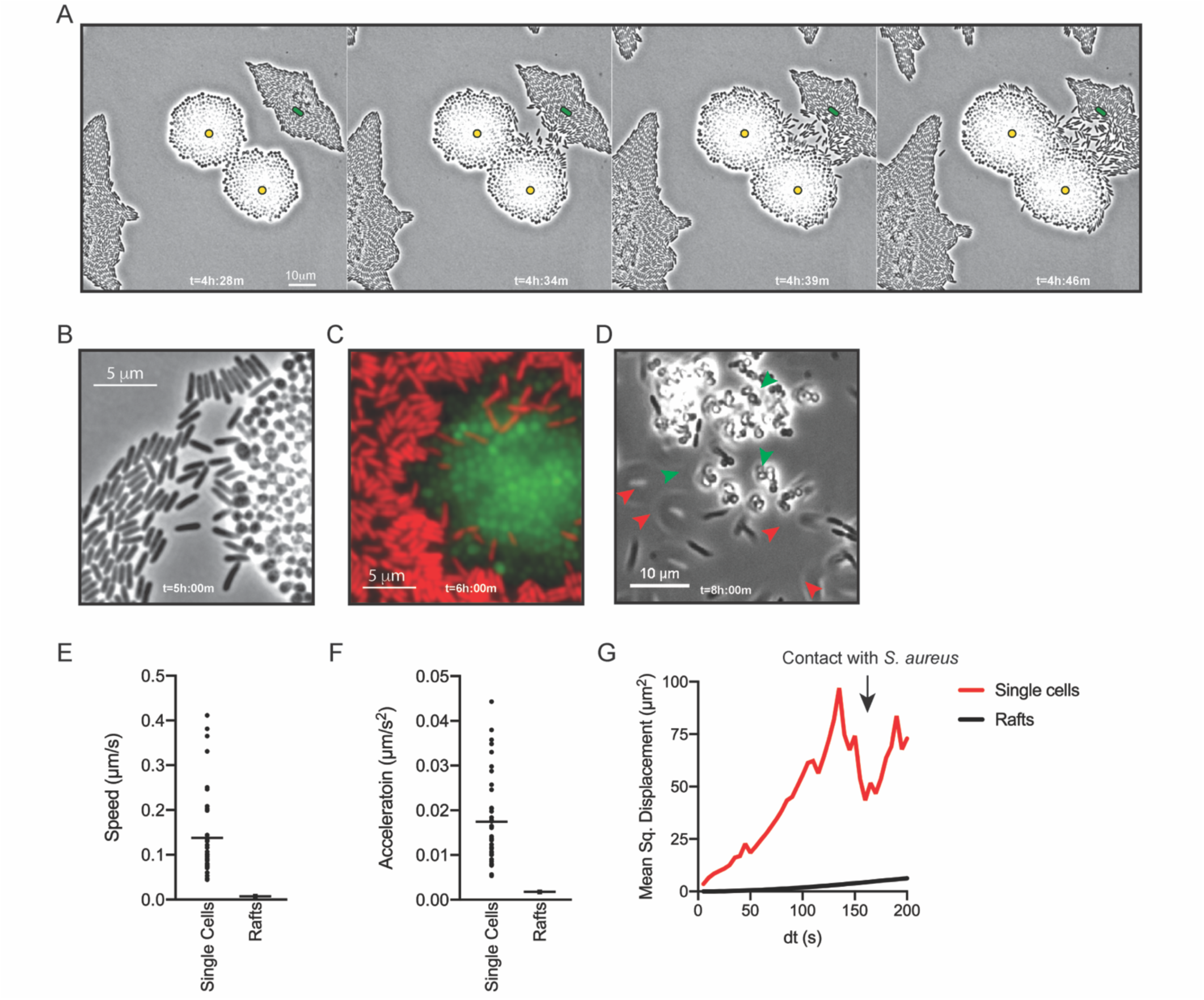
*P. aeruginosa* adopts an exploratory mode of motility in the presence of *S. aureus*. Single-cell live-imaging of *P. aeruginosa* with WT *S. aureus* (**Movie 4**). **A**. Montage of representative snap-shots are shown beginning at 4h:28m. Founding cells identified in the first frame are indicated with green rods (*P. aeruginosa*) or yellow circles (*S. aureus*). The location of the founding cell is indicated in each subsequent frame for positional reference. **B.** Snap-shot at 5h, zoomed in to visualize single cells. **C.** Snap-shot at 6h of *P. aeruginosa* (P*_A1/04/03_* -mKO) and *S. aureus* (*sarA*P1-sGFP) illustrating *P. aeruginosa* invasion into *S. aureus* colonies. **D.** Snap-shot of coculture at 8h, showing disruption of *S. aureus* colonies (green arrows) and swift-moving *P. aeruginosa* cells out of the plane of focus (red arrows). Single *P. aeruginosa* cells and the leading edge of rafts were tracked in the presence of *S. aureus* and the speed (µm/s), acceleration (µm/s^2^), and mean squared displacement (µm^2^) for four independent movies are indicated, respectively, in **E – G**.

To quantitate the movement of *P. aeruginosa* single-cell motility in comparison to collective motility in rafts, the movement of individual cells and the leading edge of the rafts was tracked over time. In comparison to cells moving in rafts, individual cells moved with increased speed (µm/s), acceleration (µm/s^2^), and mean squared displacement (MSD, µm^2^) (**Figure 2 E, F**, and **G**, respectively). MSD represents a combined measure of both the speed and directional persistence of the cell, thus an increased MSD in single cells over a change in time suggests single cells exhibit directed motion, followed by a decreased MSD when *P. aeruginosa* cells reach the *S. aureus* colony.

### *P. aeruginosa* type IV pili drive exploratory motility

How is *P. aeruginosa* motility generated in response to *S. aureus*? When grown in monoculture, *P. aeruginosa* performs cellular movement through the action of a single polar flagellum and the type IV pili (TFP) (Conrad et al. 2011; Gibiansky et al. 2010; Merritt et al. 2010). To determine how the observed *P. aeruginosa* “exploratory motility” is generated, *P. aeruginosa* strains deficient in the production of either TFP (Δ*pilA*; encoding the pilin monomers), flagella (Δ*flgK*; encoding the flagellar hook protein), or both (Δ*pilA* Δ*flgK*) were examined. Time-lapse images were taken as described for **Figure 2**, except they were acquired at 50 ms intervals for visualization of specific motility patterns. Representative movies and snap-shots (**Figure 3A**) were chosen at the time-point where *P. aeruginosa* was found to exhibit both slow and swift single-cell movements. The *pilA* mutant was unable move away from the group as single cells (**Movie 6**), as seen for the parental *P. aeruginosa* (**Movie 5**), suggesting the TFP are required for *P. aeruginosa* exploratory motility. However, the swift movement observed in the WT (**Figure 3A**, see boxed inset with red arrows), was maintained in Δ*pilA*, demonstrating that TFP are not required for this behavior.

**Figure 3.**
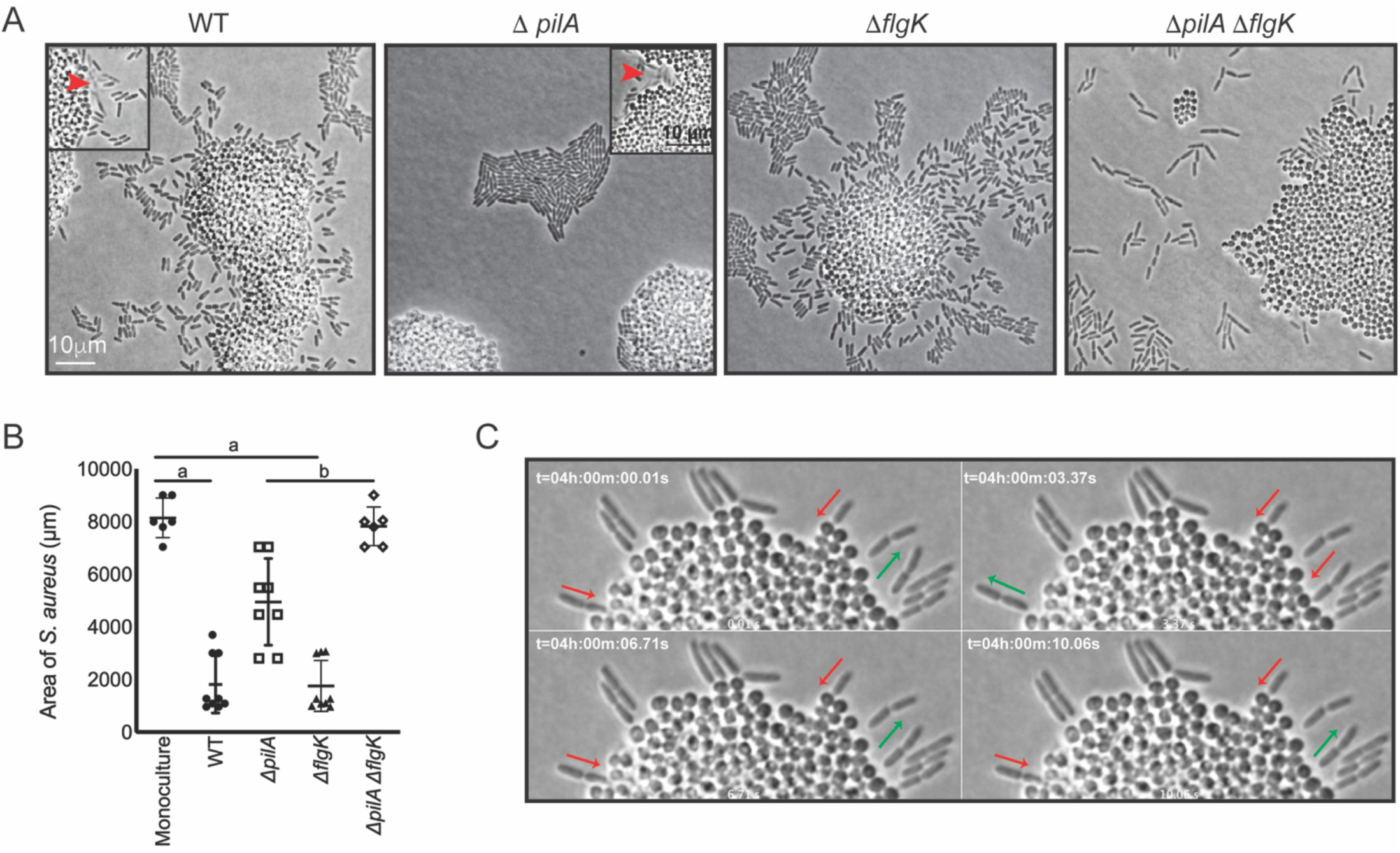
*P. aeruginosa* exploratory motility is driven by type IV pili. Live, single cell imaging of *P. aeruginosa* with WT *S. aureus*. **A**. Representative snap-shots of coculture with WT *S. aureus* and *P. aeruginosa* (WT, Δ*pilA*, Δ*flgK*, and Δ*pilA* Δ*flgK*, left to right, **Movies 5-8**, respectively) are shown at t=4.5h. Boxed insets show swift-moving *P. aeruginosa* cells out of the plane of focus (red arrows). **B.** The area of *S. aureus* in monoculture or in the presence of the indicated *P. aeruginosa* strain was calculated at t=5h by dividing the total area in a single frame by the number of *S. aureus* colonies. A minimum of four movies were analyzed per condition. The mean and standard deviation are indicated. Statistical significance was determined by one-way ANOVA followed by Tukey’s Multiple Comparisons Test - *a* indicates a statistically significant difference (*P ≤* 0.05) between *S. aureus* in monoculture and in the presence of either WT *P. aeruginosa* or Δ*flgK*; *b* indicates a statistically significant difference between Δ*pilA* and Δ*pilA* Δ*flgK*. **C.** Representative snap-shots of **Movie 5**, WT *P. aeruginosa* and *S. aureus* beginning at 4h with 50 ms intervals, showing back-and-forth motion. Red arrows indicate when a *P. aeruginosa* cell is moving in towards the *S. aureus* colony and green arrows indicate when a cell is moving away.

We next examined the Δ*flgK* mutant (**Movie 7**), which was seen to adopt the single-cell behaviors of the WT, except the swift movements were not observed, supporting the hypothesis that this movement is generated by the flagellum. These data suggest that while *S. aureus* modulates both TFP and flagella-mediated motility, the early events necessary for exploration (initiation of single cell movement and directional movement towards *S. aureus*) requires the TFP.

We also examined the response of a double Δ*pilA* Δ*flgK* mutant in the presence of *S. aureus* (**Movie 8**). Surprisingly, this mutant exhibited a phenotype distinct from either the WT, or the individual Δ*pilA* and Δ*flgK* mutants. *P. aeruginosa* cells deficient in both TFP and flagella were not only unable to produce the respective movements characteristic of these motility motors, but also were unable to remain within a raft-like group (**Figure 3A**). This behavior was not dependent upon the presence of *S. aureus*, as a similar pattern was observed for Δ*pilA* Δ*flgK* when visualized in the absence of *S. aureus* (**Figure S1**).

The influence of *P. aeruginosa* exploratory motility on *S. aureus* growth was also examined by measuring the area of the *S. aureus* colonies at 4.5 hours post coculture. *S. aureus* colonies were significantly smaller in the presence of WT *P. aeruginosa* or the Δ*flgK* mutant in comparison to *S. aureus* grown in monoculture (**Figure 3B**). However, in the presence the Δ*pilA* mutant, which was deficient in exploratory motility, colonies were significantly larger in comparison to WT and the Δ*flgK* mutant. Moreover, when grown in the presence of the double mutants, the area of the *S. aureus* colonies was not significantly different from *S. aureus* monoculture. These data suggest that ability of *P. aeruginosa* to perform exploratory motility influences *P. aeruginosa* inhibition of *S. aureus* growth.

While visualizing *P. aeruginosa*-*S. aureus* interactions at 50 ms intervals, we observed an additional *P. aeruginosa* behavior. When *P. aeruginosa* first encounters the *S. aureus* colony, the cell body orients perpendicular to the surface of the colony and moves back-and-forth (**Movie 5** and **Figure 3C**) and appears to drive *P. aeruginosa* into the *S. aureus* colony. This behavior was only observed in the WT and Δ*flgK* mutant, suggesting that it is driven by the TFP.

### Agr-regulated secreted *S. aureus* factors promote *P. aeruginosa* motility

Live-imaging suggests *P. aeruginosa* is capable of sensing *S. aureus* and initiating exploratory motility from a distance. Thus, we hypothesized that *P. aeruginosa* responds to *S. aureus* secreted factors. Since we observed that TFP were required for exploratory motility, we sought to develop a macroscopic assay where TFP motility could be monitored in the presence of *S. aureus* secreted factors only. Macroscopically, population-scale TFP-mediated twitching motility can be visualized by inoculating *P. aeruginosa* cells onto the bottom of a plate though 1.5% agar (sub-surface); cells move on the surface of the plate under the agar and can be stained with crystal violet for visualization (Turnbull and Whitchurch 2014). To test the hypothesis that *P. aeruginosa* responds to *S. aureus* secreted products, cell-free supernatant derived from overnight cultures of WT *S. aureus*, (normalized to OD_600_ = 5.0) was spread on the plate, prior to pouring molten agar. *S. aureus* supernatant significantly increased the motility diameter of *P. aeruginosa* in dose-dependent manner (**Figure 4A**), supporting the hypothesis that *S. aureus* secreted factors increase *P. aeruginosa* twitching motility.

**Figure 4.**
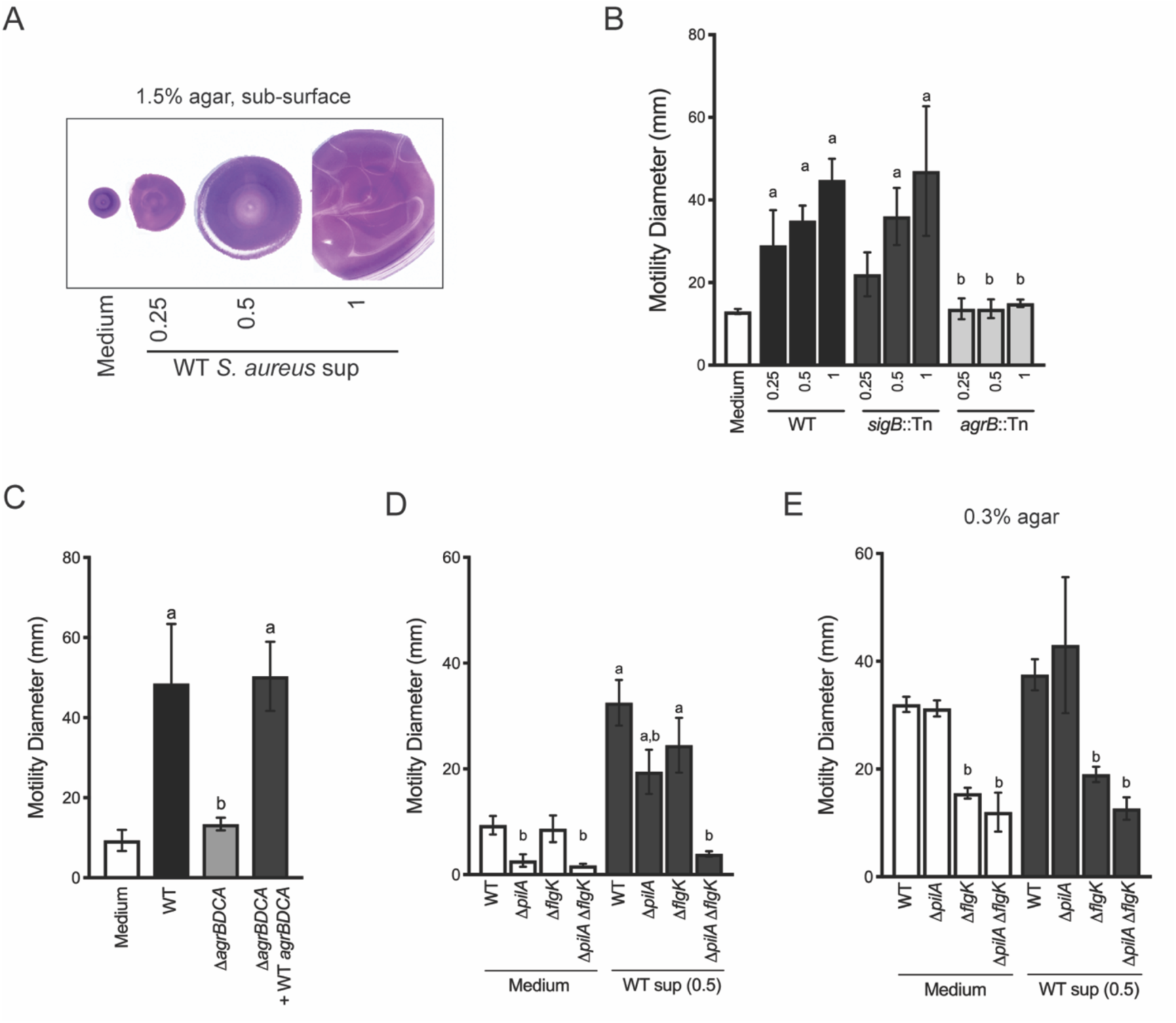
Agr-regulated secreted *S. aureus* factors increase *P. aeruginosa* motility. **A – D**. Motility of WT *P. aeruginosa* was monitored by macroscopic sub-surface inoculation assays in the presence of medium alone or cell-free supernatant from the indicated *S. aureus* strains. **A** illustrates representative motility zones stained with crystal violet for visualization. The dilution factor of the supernatant is indicated in **A** and **B**. Undilute supernatant was used in **C**. In **D** and **E**, the motility of the indicated *P. aeruginosa* mutants was analyzed in the presence of medium alone or supernatant derived from WT *S. aureus* (0.5 dilution) under 1.5% agar or within 0.3% agar (**E**). The mean and standard deviation are indicated for at least three biological replicates. Statistical significance was determined by one-way ANOVA followed by Tukey’s Multiple Comparisons Test - *a* indicates a statistically significant difference (*P ≤* 0.05) between the motility observed in the presence of *S. aureus* supernatant compared to medium alone, and *b* indicates a statistically significant difference (*P ≤* 0.05) between the motility observed in the mutant strain (*S. aureus* mutants in **B** and **C**, *P. aeruginosa* mutants in **D** and **E**) compared to the parental.

Two primary regulators of secreted factors in *S. aureus* are the alternative stress sigma factor, sigma B (SigB) (Bæk et al. 2013) and the accessory gene regulator (Agr) quorum sensing system (Nair et al. 2011). *S. aureus* strains with transposon insertions in either *sigB* or *agrB* (Fey et al. 2013) were examined for their ability to induce *P. aeruginosa* twitching motility (**Figure 4B**). While the *sigB::*Tn strain phenocopied the WT, the *agrB::*Tn mutant lost all ability to induce *P. aeruginosa* twitching motility, suggesting that Agr regulates the production of the factors promoting motility in *P. aeruginosa.* The *agr* operon is organized around two divergent promoters, P2 and P3, and generates two primary transcripts, RNAII and RNAIII, respectively. RNAII encodes AgrB, AgrD, AgrC, and AgrA (Le and Otto 2015). To confirm a role for the Agr quorum sensing system, an unmarked deletion of *agrBDCA* was generated and complemented with WT *agrBDCA. P. aeruginosa* motility was examined in the presence of supernatant derived from these strains. As predicted, the Δ*agrBDCA* mutant was unable to enhance *P. aeruginosa* motility, and complementation restored activity to WT levels (**Figure 4C**).

To confirm that the TFP are necessary for increased *P. aeruginosa* motility in this assay, we examined the response of *pilA* deficient *P. aeruginosa* to *S. aureus* supernatant. In the absence of *S. aureus* supernatant, the *pilA* mutant was required for twitching motility, as previously reported (Darzins 1994). However, while motility was reduced in the presence of *S. aureus* supernatant, the Δ*pilA* mutant retained some ability to respond to *S. aureus* secreted factors (**Figure 4D**). Since we observed during live imaging that *S. aureus* can increase *P. aeruginosa* flagella-mediated motility, in addition to TFP-mediated motility, we hypothesized that the response retained in the Δ*pilA* mutant was due to increased flagellar-mediated motility. To test this hypothesis, we examined the response of Δ*flgK* and a double Δ*pilA* Δ*flgK* mutant to *S. aureus* supernatant. The single Δ*flgK* mutant phenotype trended lower, but was not significantly different from WT, while the motility of the Δ*pilA* Δ*flgK* mutant was reduced to levels not significantly different from Δ*pilA* or Δ*pilA* Δ*flgK* in the absence of supernatant (**Figure 4D**). These data support our observation from live-imaging that, while TFP were the primary contributors to increased motility, *S. aureus* secreted factors could increase both TFP- and flagellar-mediated motility.

To formally examine the response of *P. aeruginosa* flagella-mediated motility to *S. aureus* supernatant, traditional macroscopic low-percentage agar assays that measure the contribution of flagellar motility and chemotaxis were performed. Cell-free *S. aureus* supernatant was mixed into 0.3% agar prior to inoculating *P. aeruginosa* cells into the agar. The diameter of the *P. aeruginosa* motility zone showed a modest, but not significant increase in the presence of *S. aureus* supernatant, compared to medium alone (**Figure 4E**). To examine the role for flagella in response to *S. aureus* under swim assay conditions, the *flgK* mutant was tested. As expected, flagellar-mediated motility, both with and without *S. aureus* supernatant, was significantly reduced. To determine if TFP contribute under these assay conditions, the Δ*pilA* and Δ*pilA* Δ*flgK* mutants were also examined. The motility diameter of Δ*pilA* was not significantly different from WT, and the double mutant phenocopied the single *flgK* mutant, suggesting that flagella are primarily responsible for the motility observed here.

### *P. aeruginosa* biases the directionality of movement up a concentration gradient of *S. aureus-*secreted factors

Plate-based macroscopic motility assays measure both an absolute increase in motility and directional movement up a self-generated gradient (chemotaxis) as bacterial populations metabolize the available substrates and expand outward radially (Shapiro 1984). While flagella-based movement though liquid has been extensively described in *P.* aeruginosa for a variety of chemoattractants, directional movement on a surface is poorly understood. Kearns and Shimkets previously reported *P. aeruginosa* pili-mediated biased movement up a gradient of phosphatidylethanolamine (PE) on the surface of an agar plate (Kearns, Robinson, and Shimkets 2001). To determine if *P. aeruginosa* is capable of moving up a previously established concentration gradient of *S. aureus* supernatant, supernatant derived from either WT or the Δ*agrBDCA* mutant of *S. aureus* was spotted onto the surface of 1.5% agar (containing medium buffered for pH only, no carbon source). The secreted factors were allowed to diffuse and establish a gradient for approximately 24 hours, prior to inoculating *P. aeruginosa* onto the surface, as previously described (Kearns, Robinson, and Shimkets 2001). Preferential movement of *P. aeruginosa* towards supernatant derived from WT *S. aureus* was observed, but not for medium alone or supernatant derived from the Δ*agrBDCA* mutant (**Figure 5A**). The response for the Δ*pilA* mutant was also examined and all motility was abrogated in this mutant, demonstrating motility observed in this assay is entirely pili-mediated. These data support the hypothesis that *P. aeruginosa* biases the directionality of TFP-mediated motility up a concentration gradient of *S. aureus* secreted factors.

**Figure 5.**
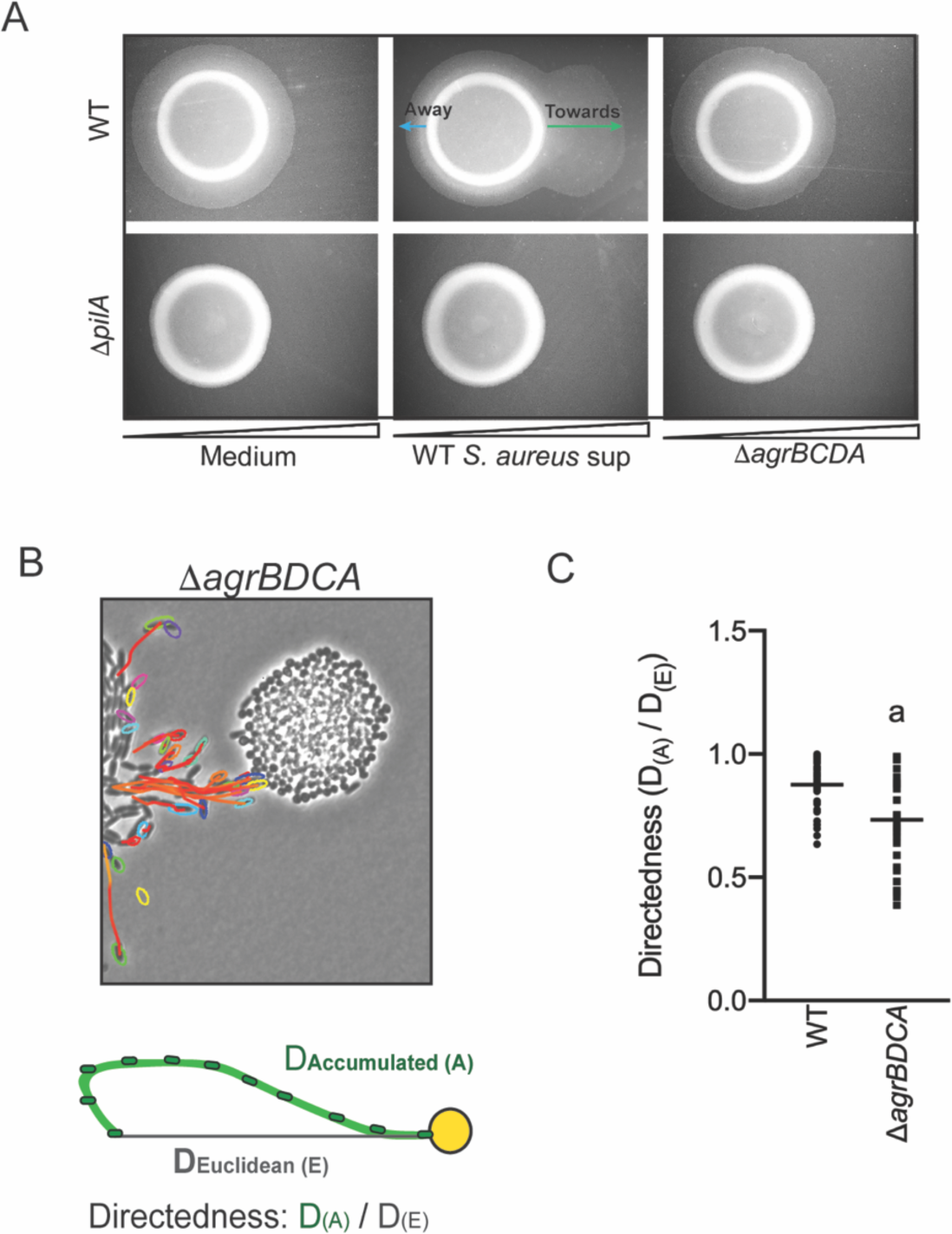
*P. aeruginosa* biases the directionality of movement up a concentration gradient of Agr-regulated secreted factors. **A.** A concentration gradient of either *S. aureus* growth medium (TSB), *S. aureus* supernatant derived from WT or Δ*agrBCDA* was established by spotting onto the surface of the agar and allowing a concentration gradient to establish by diffusion for approximately 24h, prior to spotting *P. aeruginosa* onto the agar (6 mm to the left) and surface-based motility imaged after 24h. Representative images of at least three independent experiments are shown. **B**. Example of live imaging of WT *P. aeruginosa* with *S. aureus* Δ*agrBCDA* with tracks of single cells shown and schematic illustrating the methods for calculating the directedness. Single *P. aeruginosa* cells were tracked from first frame a cell exited the raft to the frame where it first encounters *S. aureus*. The accumulated track distance, D_(A)_, was measured for at least 30 cells in four independent movies and compared to the Euclidean distance, D_(E)_, between the position of the cell in the first and last frame tracked. The ratio of D_(A)_ / D_(E)_ (Directedness) is shown in **C** for *P. aeruginosa* towards WT *S. aureus* compared to Δ*agrBDCA* with the mean indicated. Statistical significance was determined by an unpaired Student’s *t*-test (*a* = *P ≤* 0.05).

We next asked if *P. aeruginosa* would also fail to migrate towards *S. aureus* deficient in Agr activity at the single cell level. WT *P. aeruginosa* was visualized in coculture with Δ*agrBDCA*, as previously performed in the presence of WT *S. aureus* – with images acquired every 5s for 8 hours (**Movie 4 for WT: Figure 2A** and **Movie 9 for the Δ*agrBDCA* mutant: Figure 5B**). In comparison to WT *S. aureus, P. aeruginosa* behavior was significantly altered in the presence of Δ*agrBDCA. P. aeruginosa* remained capable of initiating single cell movement; however, once cells initiated single-cell movement, the path of their movement did not appear as directed towards the *S. aureus* colonies. In fact, some cells seemed to actively avoid the *S. aureus* colony all together.

To quantify the directedness of *P. aeruginosa* movement in the presence of WT and the Δ*agrBDCA*, single *P. aeruginosa* cells were tracked from the first frame a cell exited the raft to the frame where it first encounters *S. aureus* (**Figure 5B**). The accumulated track distance, D_(A)_, was measured and compared to the Euclidean distance, D_(E)_, between the position of the cell in the first and last frame tracked. The directness of *P. aeruginosa* towards *S. aureus* was calculated as a ratio of D_(A)_ / D_(E)_. *P. aeruginosa* exhibited a higher directedness ratio towards WT *S. aureus* compared to the Δ*agrBDCA* mutant (**Figure 5C**). These data support the hypothesis that Agr regulates the production of *S. aureus* secreted factors driving the directionality of *P. aeruginosa* motility.

### *P. aeruginosa* responds to *S. aureus* with a decrease in cAMP levels

How does *P. aeruginosa* sense the presence of *S. aureus* secreted products and initiate TFP-driven exploratory motility? *P. aeruginosa* encodes three known chemotaxis pathways: two flagella-mediated (*che* and *che2*) and a putative *TFP*-mediated system (*pil-chp*) (Darzins 1994; Whitchurch et al. 2004). While several proteins encoded in the Pil-Chp pathway comprise a signal transduction pathway very similar (by gene homology) to the flagella-mediated chemotaxis pathway, their role in directional twitching motility remains unclear. A significant challenge lies in that many of these proteins are required for pilus assembly, thus teasing apart their requirement for motility *per se* verses regulation of a chemotactic response is challenging. Nonetheless, *pilJ* is predicted to encode the TFP methyl-accepting chemoreceptor protein (MCP) and by homology to the flagella MCPs, PilJ is expected to detect changes in the concentration of an attractant or repellent and initiate a signaling cascade to control twitching motility.

To determine if PilJ is necessary for *P. aeruginosa* to sense *S. aureus*, we examined the response of a Δ*pilJ* mutant to *S. aureus* (WT) by single-cell imaging. In the presence of *S. aureus*, the Δ*pilJ* mutant phenocopied the behavior of WT *P. aeruginosa*; moving as single cells and with preferred directionality towards *S. aureus* (**Figure 6A, left**), suggesting that PilJ is not necessary for exploratory motility. Surprisingly however, when Δ*pilJ* was imaged in the absence of *S. aureus*, this strain was capable of adopting many of the behaviors we observed in the presence of *S. aureus* (i.e. increased single-cell movement, **Figure 6A center, Movie 10**).

**Figure 6.**
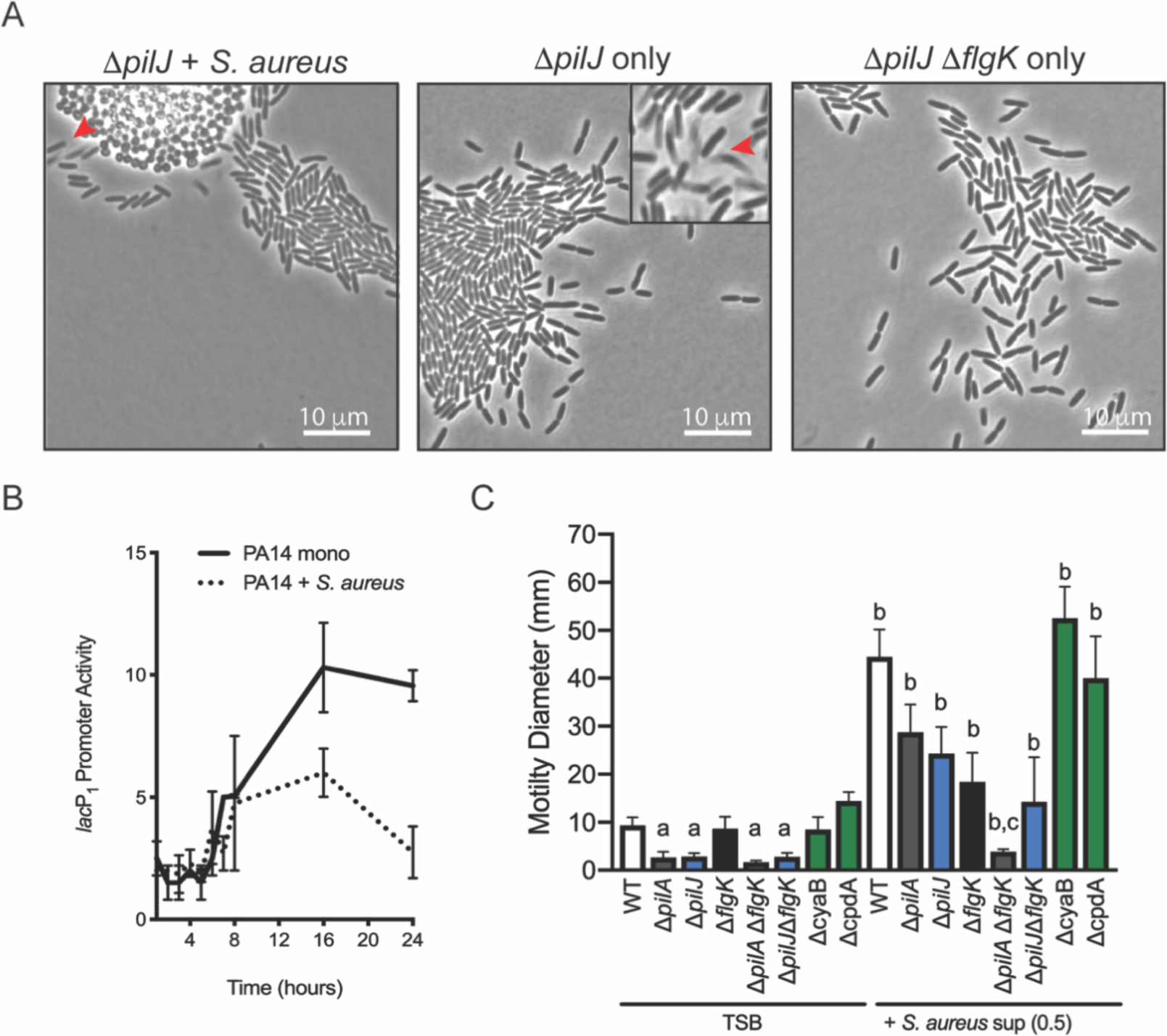
*P. aeruginosa* exploratory motility is driven by modulation of cAMP levels. **A**. Representative snap-shots at t=4h of live imaging of *P. aeruginosa* Δ*pilJ* with WT *S. aureus* (left), Δ*pilJ* alone (**Movie 10**, center), and Δ*pilJ* Δ*flgK* alone. Red arrows indicate swimming cells. **B**. Kinetic analysis of cAMP levels in *P. aeruginosa* cells grown on an agar surface in monoculture and with *S. aureus.* The cellular cAMP level is expressed as *lac*P1*-lacZ* activity divided by the vector control. The mean +/- standard deviation of three biological replicates is indicated. **C.** Macroscopic twitch assay performed as previously described for the indicated mutants. The mean and standard deviation are indicated for at least three biological replicates. Statistical significance was determined by one-way ANOVA followed by Tukey’s Multiple Comparisons Test - *a* indicates a statistically significant difference (*P ≤* 0.05) between the motility observed in the mutant strain compared to the parental, *b* indicates a statistically significant difference (*P ≤* 0.05) in motility observed in the presence of *S. aureus* supernatant compared to medium alone, and *c* indicates a statistically significant difference (*P ≤* 0.05) between Δ*flgK* and Δ*flgK* Δ*pilA* in the presence of supernatant.

How might sensing *S. aureus* and eliminating PilJ have similar phenotypic outcomes for *P. aeruginosa*? We previously reported that PilJ participates in a hierarchical regulatory cascade of the second messengers, cyclic AMP (cAMP) and cyclic di-GMP (c-di-GMP), which regulate transitions between motility and surface attachment in *P. aeruginosa*. In the absence of PilJ, cells are predicted to have low intracellular levels of cAMP and c-di-GMP, with a concomitant decrease in surface attachment and increase in flagella-mediated motility (Luo et al. 2015). Indeed, some of the movement in the *pilJ* mutant had visual appearance attributed in flagella-mediated motility in **Figure 2** (red arrows in **Figure 6A**). To determine if increased flagella mediated-motility is responsible for the phenotypes observed during live imaging of the Δ*pilJ* mutant, we constructed a Δ*pilJ* Δ*flgK* double mutant. While the double mutant had significantly reducing swimming motility in the macroscopic motility assay (**Figure S2**), its phenotype in the live, single-cell imaging remained similar to both WT *P. aeruginosa* with *S. aureus* and Δ*pilJ* alone, suggesting that the Δ*pilJ* mutant is not only able to modulate TFP-mediated motility at the single cell level, but also adopts characteristics of exploratory motility.

Since PilJ is necessary for *P. aeruginosa* to increase cAMP in response to contact with a surface, we hypothesized that *S. aureus* might influence *P. aeruginosa* cAMP levels. To test this model, we utilized a cAMP-dependent promoter fusion, *lac*P1*-lacZ*, whose activity has been shown to reflect cellular cAMP levels (Fulcher et al. 2010). We deduced cellular cAMP levels based on the fold change of *lac*P1*-lacZ* activity over a vector control, as reported (Fulcher et al. 2010; Luo et al. 2015), and measured these levels over time in the presence of *S. aureus*, while grown on the surface of an agar plate. For the first eight hours of coculture, *P. aeruginosa* cAMP levels in the presence of *S. aureus* were indistinguishable from *P. aeruginosa* grown in monoculture (**Figure 6B**). However, after eight hours, cAMP levels continue to rise when *P. aeruginosa* is grown in monoculture, as previously reported (Luo et al. 2015), but when cocultured with *S. aureus*, a similar increase in cAMP was not observed. Although the conditions and kinetics of this bulk cell assay are delayed from the single-cell live imaging, these data support the hypothesis that *P. aeruginosa* responds to *S. aureus* by modulating cAMP levels.

To further examine a role for *P. aeruginosa* cAMP modulation in response to *S. aureus*, we examined the ability of the Δ*pilJ* mutant to respond to *S. aureus* supernatant in the macroscopic twitching motility assay. As previously reported (Fulcher et al. 2010), a Δ*pilJ* mutant is deficient in macroscopic twitching motility; however, consistent with the live imaging, the Δ*pilJ* mutant remained capable of responding to *S. aureus* supernatant (**Figure 6C**). We also tested the double Δ*pilJ* Δ*flgK* mutant, which demonstrated a reduced response, in comparison to the single Δ*pilJ* mutant. However, it maintained the ability to respond to *S. aureus*, in comparison to the double Δ*pilA* Δ*flgK* mutant. These data support a model whereby the Δ*pilJ* mutant maintains the capacity to respond to *S. aureus* and to undergo TFP-mediated motility. To formally examine the role for cAMP, we tested a mutant in the *cyaB* gene, encoding the primary adenylate cyclase required for the generation of cAMP and a mutant in the *cpdA* gene, encoding the phosphodiesterase required to convert cAMP into ATP (Fulcher et al. 2010). Consistent with our model, Δ*cyaB* (low cAMP), demonstrated an enhanced response to *S. aureus* (**Figure 6C**). However, the phosphodiesterase mutant, predicted to have high cAMP (Δ*cpdA*), phenocopied the WT. These data support a model whereby *S. aureus* reduces cellular cAMP at the level adenylate cyclase and not though activation of CpdA (see Model, **Figure 8** and discussion).

**Figure 7.**
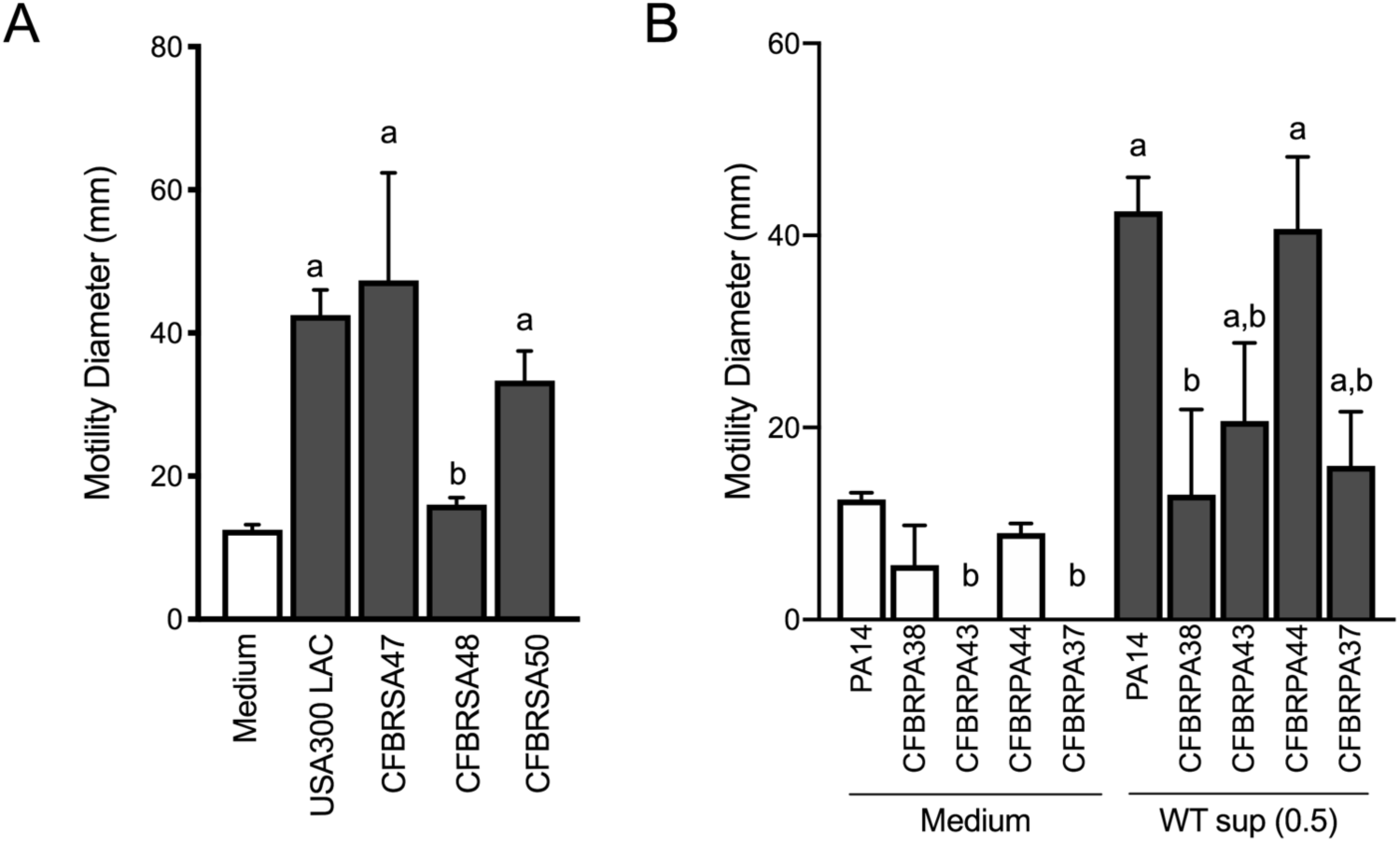
*P. aeruginosa* and *S. aureus* isolates from CF patients participate in interspecies signaling. **A**. The motility of laboratory strain *P. aeruginosa* stain PA14 was monitored in the presence of undilute cell-free supernatant from clinical *S. aureus* CF isolates, CFBRSA47, 48, and 50, in comparison to the positive control *S. aureus* strain USA300 LAC. **B**. The motility of clinical *P. aeruginosa* isolates (CFBRPA38, 43, 44, and 37), in the presence of cell-free supernatant derived from *S. aureus* USA300 LAC (0.5 dilution) is indicated, in comparison to *P. aeruginosa* PA14. The mean and standard deviation are indicated for at least three biological replicates. Statistical significance was determined by one-way ANOVA followed by Tukey’s Multiple Comparisons Test - *a* indicates a statistically significant difference (*P ≤* 0.05) between the motility observed in the presence of *S. aureus* supernatant compared to medium alone, and *b* indicates a statistically significant difference (*P ≤* 0.05) between the motility observed in the CF isolates, in comparison laboratory strains (*S. aureus* USA300 LAC in **A** and *P. aeruginosa* laboratory strain in **B**).

**Figure 8.**
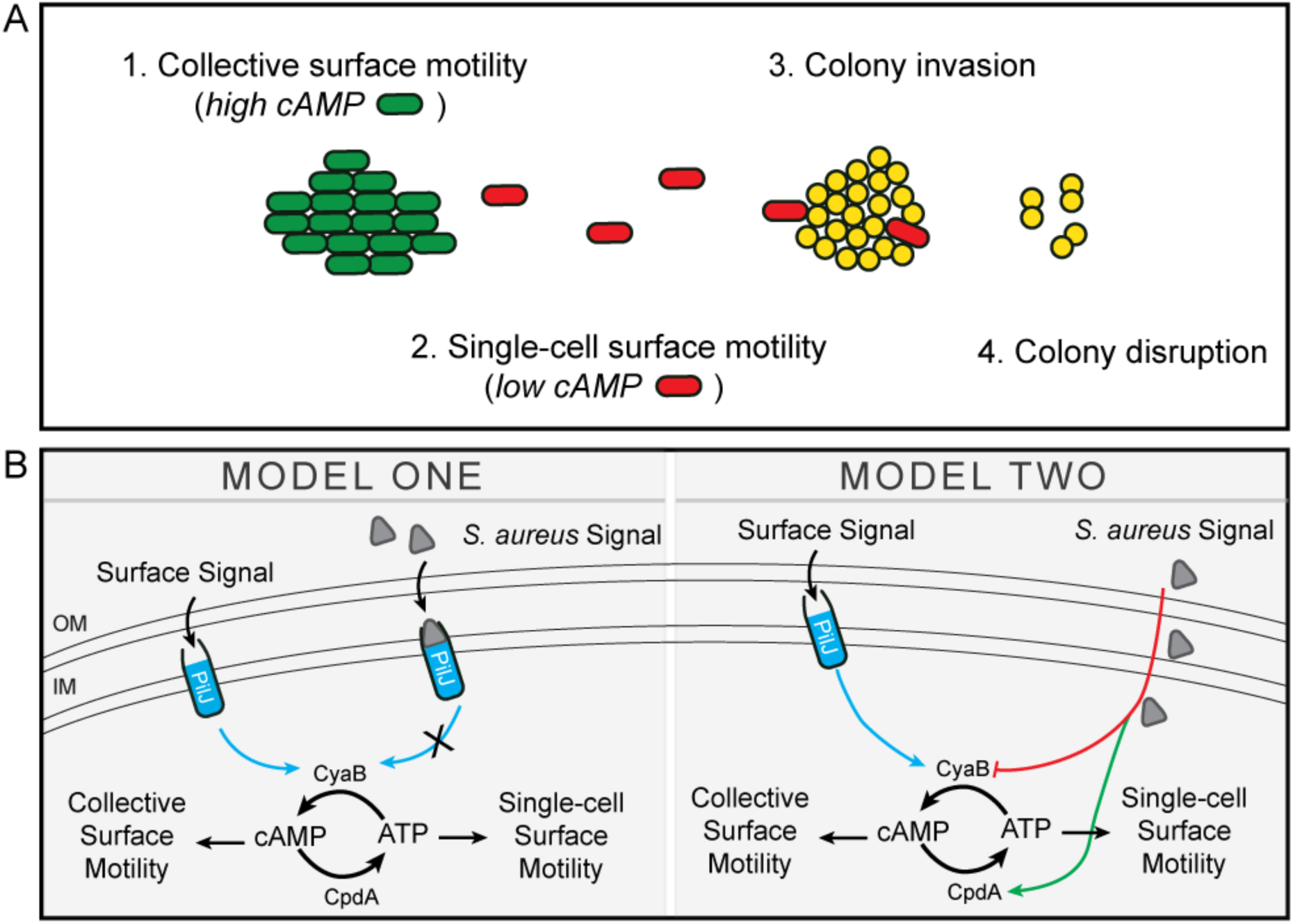
Proposed model of *S. aureus* induced *P. aeruginosa* exploratory motility. **A.** In monoculture *P. aeruginosa* minimally traverses surfaces via collective TFP-mediated motility in large rafts, modulated in part, through increased levels of the second messenger, cAMP. When *P. aeruginosa* senses the presence of *S. aureus*, single cells exit the group (with an associated decrease in cAMP), move towards *S. aureus*, invade, and disrupt the colony. **B.** Two proposed models for how *S. aureus* signals modulate cAMP. In Model One, *S. aureus* signals bind to the chemoreceptor, PilJ and inhibits its ability to activate the adenylate cyclase, CyaB, resulting in decreased cAMP levels. In Model Two, *S. aureus* signals lower cAMP levels through an unknown, PilJ-independent mechanism by either decreasing activity of the adenylate cyclase, CyaB or increasing the activity of the phosphodiesterase, CpdA (or both).

### Clinical isolates of *P. aeruginosa* and *S. aureus* perform exploratory motility

Our data thus far demonstrate the *P. aeruginosa* can sense *S. aureus* from a distance and modulate both flagella and TFP-mediated motility. To begin to examine if these behaviors might be relevant during airway infection in CF patients, we sought to determine if *S. aureus* isolates from CF patients are capable of inducing *P. aeruginosa* motility in the macroscopic twitching assay. Supernatant derived from two out of three clinical CF *S. aureus* isolates (each from different patients) was capable of inducing motility of the WT *P. aeruginosa* laboratory strain, to an extent not significantly different from that previously observed with WT *S. aureus* (**Figure 7A**). Since we previously observed Agr was necessary for *S. aureus* to promote motility, we hypothesized that CFBRSA48 was unable to induce *P. aeruginosa* due to reduced activity of the Agr quorum sensing system – an adaptation previously reported for *S. aureus* CF isolates (Nair et al. 2011; Goerke and Wolz 2010). Since Agr also positively regulates the production of *S. aureus* α-hemolysin, hemolysis on sheep blood agar plates is often used as an indicator of Agr activity (Peng et al. 1988). However, all three clinical isolates maintained WT levels of hemolysis, suggesting Agr is active in these strains and an unknown mechanism accounts for reduced activity towards *S. aureus* in CFBRSA isolate 48 (**Figure S3**).

Next, we examined the capacity of a panel of *P. aeruginosa* isolates from CF patients to respond to *S. aureus*. Phenotypic loss of twitching motility in monoculture is often observed in chronic *P. aeruginosa* isolates from CF patients (Mayer-Hamblett et al. 2014), thus it was not surprising that two of the four isolates examined exhibited no detectable twitching motility in the absence of *S. aureus*. Nonetheless, each *P. aeruginosa* isolate was able to respond to *S. aureus* supernatant, although to varying degrees (**Figure 7B)**. Importantly, even mucoid *P. aeruginosa* isolates (38, 43, 37), exhibited at least a two-fold increase in motility in the presence of *S. aureus* supernatant, above medium alone.

## Discussion

Current massive sequencing efforts yield unprecedented information regarding microbial community composition during infection, but fail to provide the necessary information required to functionally understand microbial behaviors in mixed species communities. Moreover, while the bulk assays most frequently utilized to study microbial interactions have provided insight into how bacterial species can influence community survival, metabolism, or virulence factor production (Hotterbeekx et al. 2017), we lack detailed information regarding how a single cell responds to the presence of another species. Here, we developed quantitative live-imaging methods to visualize the behaviors of two clinically important organisms and track their behavior overtime. Through these studies, we uncovered microbial behaviors that could not have been predicted from bulk assays alone. We observed that *P. aeruginosa* is able to sense *S. aureus* from a distance by responding to secreted factors. *P. aeruginosa* is then able to tune its motility patterns, adopting a behavior we refer to as ‘exploratory motility’. The studies presented here open many questions regarding how *P. aeruginosa* is able to sense and respond to other microbial species. Our current working model is illustrated in **Figure 8** and some of the many questions produced from these studies are discussed in detail below.

What interspecies signals does *P. aeruginosa* sense and respond to? Our data thus far reveal that *P. aeruginosa* can sense secreted products from *S. aureus*. We also show that these products are regulated by the Agr quorum sensing system in *S. aureus.* Identification of the nature of these interspecies signals, as well as investigation into how conserved these processes are among other microbial species is underway. Our current data suggest that *P. aeruginosa* can sense multiple secreted factors from *S. aureus.* This hypothesis is supported by the observation that an Agr-deficient strain is unable to induce motility in the macroscopic motility assays, but remains capable of promoting single cell motility during live imaging. Thus, it is formally possible that separate signals are required for initiating single-cell motility and directional motion. Live imaging also suggests that *P. aeruginosa* cells may actively avoid Agr-deficient *S. aureus*, raising the possibility that the Agr system negatively regulates a secreted factor that is unfavorable for *P. aeruginosa*.

When *P. aeruginosa* senses *S. aureus* signals, we observed a transition from collective TFP-mediated movement to single-cell motility. TFP-mediated motility is most often described as a behavior where cells are primarily motile only in large groups. The behaviors for WT *P. aeruginosa* in monoculture observed here are consistent with previous descriptions of twitching motility, including the movement of cells in rafts, preferentially aligned along their long axis, and group tendril formation at the edges of the expanding community (Burrows 2012). In contrast, in the presence of *S. aureus, P. aeruginosa* cells were seen to exit an expanding raft and transition to single-cell motility. In the process of interrogating *P. aeruginosa* genes necessary for response to *S. aureus*, we also uncovered pathways which were necessary for *P. aeruginosa* transition between collective and individual movements, in the absence of *S. aureus*. First, while a *P. aeruginosa* mutant deficient in TFP was unable to exhibit characteristics of twitching motility, the cells also exclusively remained tethered to the group while dividing and expanding outward. However, when flagella were also deleted from this strain, the community appeared to lose its integrity and fall apart. We had initially predicted that maintenance of cells within a group under these conditions (cells sandwiched between a 2% agarose pad and a coverslip) resulted from immobilization of the cells under the pad, preventing *P. aeruginosa* from generating sufficient force to become motile by either the TFP or flagellum to move away from the group. However, the observation that the Δ*pilA* Δ*flgK* mutant was unable to retain group structure suggests the flagella may be important to actively maintain cell-cell contact during collective movement on a surface.

Additional insight into *P. aeruginosa* regulation of collective verses single cell behaviors was revealed through investigation of a *P. aeruginosa* strain deficient in PilJ. PilJ is a predicted chemoreceptor, by gene homology to the flagella chemotaxis system (Darzins 1994); however, its functional role in chemotaxis remains unclear. Instead, PilJ participates in sensing surfaces and relaying this information to the cell via modulation of cAMP (Luo et al. 2015; Persat et al. 2015). PilJ is required for twitching motility in macroscopic twitching motility assays (Fulcher et al. 2010; Kearns, Robinson, and Shimkets 2001); a phenotype we confirmed here in monoculture. However, in both our macro- and microscopic assays, the Δ*pilJ* mutant maintained the ability to undergo TFP-mediated motility in the presence of *S. aureus.* In fact, the Δ*pilJ* mutant was visualized to retain motility patterns characteristic of TFP-mediated motility in monoculture during live-imaging. Together these data suggest that the Δ*pilJ* mutant is indeed capable of performing TFP-mediated motility, but fails to expand outward sufficient to generate a macroscopic motility zone under standard macroscopic conditions – a phenotype characteristic of classic chemotaxis systems.

We also observed that similar to the Δ*pilA* Δ*flgK* mutant, the Δ*pilJ* mutant failed to remain as a group during live imaging, but instead initiated single cell movement. However, once the *pilJ* mutant (and Δ*pilJ* Δ*flgK*), exited the raft, the cells traveled with increased speed and directional persistence, in comparison to the Δ*pilA* Δ*flgK* mutant whose single cells were unable to remain tethered, but did not significantly move away from their initial location. The Δ*pilJ* mutant produces reduced levels of the second messengers cAMP and c-di-GMP, resulting in increased swarming motility (Luo et al. 2015), raising the possibility that modulation of the flagellum may influence initiation of single-cell motility in this strain as well, although further studies are necessary to examine this hypothesis.

The observation that *S. aureus* reduces *P. aeruginosa* cAMP levels further supports the hypothesis that modulation of cAMP is necessary for exploratory motility. However, whether maintenance of low cAMP levels in the presence of *S. aureus* drives the initiation of single-cell motility on a surface or is concurrent with, is unknown. The observation that the PilJ-deficient *P. aeruginosa*, which also exhibits low cAMP levels, responds similarly to *S. aureus* as WT and in monoculture also travels as single cells, leads us to posit two models. First, it is possible that *S. aureus* secreted factors bind to PilJ and inhibit its ability to increase cAMP levels. Second, *S. aureus* factors inhibit cAMP production independent of PilJ. In this model, *S. aureus* could lower cAMP levels by inhibiting CyaA/B or by activating the phosphodiesterase, CpdA. Studies to differentiate these hypotheses are underway (**Figure 8**).

*Myxococcus xanthus* is one of the most notorious predatory bacteria – elaborating an array of social behaviors reminiscent of what we observe here for *P. aeruginosa. M. xanthus* coordinates a cooperative, density-dependent feeding behavior, resulting in propulsion of the cells rapidly through the colony of prey, leading to prey lysis, and nutrient acquisition. When *M. xanthus* contacts prey cells, pili-dependent reversals are stimulated, which keeps the cells “trapped” near the prey colony, promoting contact-dependent prey killing and increased local concentration of antimicrobials (Muñoz-Dorado et al. 2016). Similarly, we observed that when *P. aeruginosa* encounters *S. aureus*, the *P. aeruginosa* cells appear to mount a coordinated response whereby *P. aeruginosa* surrounds the *S. aureus* colony and eventually invades the colony – a behavior referred to as the “wolf pack” strategy. *P. aeruginosa* produces an arsenal of secreted antimicrobials shown to inhibit the growth of *S. aureus*, yet these secreted factors alone are insufficient for cellular lysis (Limoli and Hoffman 2019). Thus, it is interesting to speculate that these behaviors function to synergistically increase cellular contacts necessary to kill *S. aureus* by a contact-dependent mechanism and to locally increase the concentration of secreted products.

*P. aeruginosa* and *S. aureus* can be coisolated from the lungs of approximately 30% of patients with CF, which is associated with poor clinical outcomes (Limoli et al. 2016). Whether interactions between these pathogens during infection directly influence pulmonary decline is unknown. However, in addition to regulation of surface behaviors, cAMP is a primary regulator of *P. aeruginosa* virulence factors (Wolfgang et al. 2003), which may influence pathogenic behaviors for these bacteria during polymicrobial infection. Moreover, during chronic infection we find *P. aeruginosa* evolves phenotypes more permissive to growth with *S. aureus* (Limoli et al. 2017), suggesting a potential advantage for *P. aeruginosa* to interact with *S. aureus* during infection.

By acquiring a fundamental understanding of how bacteria sense and respond to life with each other, we move closer to learning how to rationally manipulate interspecies behaviors during infection and in the environment. While for CF patients, this may mean preventing *P. aeruginosa* and *S. aureus* physical interactions, in other instances, we might bring species together who synergize to produce a beneficial compound.

## Materials and Methods

### Bacterial strains and culture conditions

*P. aeruginosa* and *E. coli* were routinely cultured in lysogeny broth (LB; 1% tryptone, 0.5% yeast extract, 1% sodium chloride) and *S. aureus* in tryptic soy broth (TSB, Becton Dickenson) at 37°C, with aeration. For coculture assays, both species were grown in TSB or M8 minimal medium (48 mM sodium phosphate dibasic, 22 mM potassium phosphate monobasic, 8.6 mM sodium chloride, 2.0 mM magnesium sulfate, 0.1 mM calcium chloride) supplemented with 1% glucose and tryptone. When necessary for strain construction, medium was supplemented with the following antibiotics: gentamicin (10 µg/ml *E. coli*; 30 µg/ml *P. aeruginosa*), ampicillin (100 µg/ml *E. coli*), carbenicillin (200 µg/ml *P. aeruginosa*), or chloramphenicol (10 µg/ml *S. aureus*).

*S. aureus genetic manipulation.* In-frame deletion of the *agrBDCA* operon in a *S. aureus* USA300 LAC strain (JE2) was generated using pMAD-mediated allelic replacement (Arnaud et al. 2004). Briefly, a pMAD deletion vector was created by Gibson assembly (Gibson et al. 2009) using EcoRI/BamHI-linearized pMAD and PCR amplicons of 1kb regions up- and downstream of *agrBDCA* (primer sets agr-a/agr-b and agr-c/agr-d, respectively). The vector to chromosomally complement this deletion strain was similarly produced using a Gibson assembly on the EcoRI/BamHI-lineararized pMAD vector and the PCR product of the agr-a/agr-d primers. Following the standard protocol of heat shift on selective media, strains with successful deletions and restorations of *agrBDCA* were verified by PCR analysis and chromosomal DNA sequencing.

*P. aeruginosa genetic manipulation.* In-frame deletion mutants in *P. aeruginosa* were constructed via allelic exchange as previously described (Shanks et al. 2006). DNA fragments were amplified from *P. aeruginosa* PA14 genomic DNA by PCR to generate upstream and downstream fragments of the *flgK* gene with nucleotide tails complementary to plasmid pMQ30 using the following primer pairs: flgK KO P1/flgK KO P2 and flgK KO P3/flgK KO. PCR products were cloned into pMQ30 by *in vivo* homologous recombination in *Saccharomyces cerevisiae* INV*Sc*1 (Invitrogen) as previously described (Shanks et al. 2006). The pMQ30-*flgK* deletion construct was transformed by electroporation into *E. coli* strain S17 and introduced into *P. aeruginosa* Δ*pilJ* (SMC2992) mutants by conjugation. Integrants were isolated on Vogel-Bonner minimal medium (VBMM; 10 mM sodium citrate tribasic, 9.5 mM citric acid, 57 mM potassium phosphate dibasic, 17 mM sodium ammonium phosphate, 1 mM magnesium sulfate, 0.1 mM cal-cium chloride, pH 7.0) agar with gentamicin (30 µg/ml) followed by sucrose counterselection. Resolved integrants were confirmed by PCR and sequencing.

### Live imaging of interspecies interactions

Bacteria were inoculated between a coverslip and an agarose pad and imaged with time-lapse microscopy. Pads were made by adding 900 µl of M8 medium with 1% glucose, 1% tryptone, and 2% molten agarose to a 10 mm diameter silicone mold on a coverslip. A second coverslip was placed on top of the agarose while still molten. Pads were allowed to dry for 1h at room temperature, followed by 30m at 37°C. Meanwhile, bacteria were prepared by growing to mid-log phase in M8 medium with 1% glucose and 1% tryptone, diluting to OD_600_ = 0.15 in warm media, and mixing *P. aeruginosa* and *S. aureus* 1:1. 3 µl of inoculum was spotted evenly onto the center of a warm 35 mm glass bottom dish, #1.5 mm coverglass (Cellvis). The agarose pad was removed from the mold and placed on top of the bacterial inoculum. Bacteria were immediately imaged with an inverted Nikon TiE or Ti2 at 37°C for 8h. Images were acquired with a 100x oil objective (1.45NA) with phase contrast and an ORCA Flash4.0 Digital CMOS camera (Hammamatsu).

Movies were generated and cells tracked and analyzed in Nikon Elements with General Analysis 3. For single cells, binary images were generated and single *P. aeruginosa* cells were identified and tracking began when the first *P. aeruginosa* cell exited a raft. Rafts were tracked up until the time point where single cell tracking began. Rafts edges were manually identified. The speed (µm/s), acceleration (µm/s^2^), mean squared displacement (µm^2^), accumulated track distance, D_(A)_, and the Euclidean distance, D_(E)_, were measure for at least 30 tracks (for single cells) in four independent movies. The area of *S. aureus* was determined by dividing the total area occupied in a field of view divided by the number of colonies.

### Macroscopic Coculture Assays

*P. aeruginosa* sub-surface twitch (Turnbull and Whitchurch 2014) and 0.3% soft agar motility assays (Ha, Kuchma, and O’Toole 2014) were performed as previously described with modifications for treatment with *S. aureus* supernatant. *S. aureus* was grown in TSB to OD_600_ = 5.0, centrifuged to pellet cells, and supernatant filtered through a 0.22 µm filter. For soft agar assays, cell-free supernatant was mixed at the indicated concentrations with cooled, but still molten agar prior to pouring. For twitch assays, 100 µl of supernatant was spread onto the bottom of a petri plate before pouring media (1.5% agar). *P. aeruginosa* was grown to OD_600_ = 5.0 in TSB and inoculated into motility plates by either stabbing with a toothpick halfway through the agar (0.3%), all the way though (sub-surface). Plates were incubated at 37°C for 24h, followed by 24h at room temperature. The diameter of the motility zones was measured in mm. For twitch plates, agar was dropped out and the *P. aeruginosa* biomass stained with 1% crystal violet for visualization.

### Directional Twitching Motility Assay

Experiments were performed as previously described (R. M. Miller et al. 2008). In brief, buffered agar plates (10 mM Tris, pH 7.6; 8 mM MgSO_4_; 1 mM NaPO_4_, pH 7.6; and 1.5% agar) were poured and dried at room temperature for 24h. 4 µl of supernatant or medium (TSB) was spotted on the plates, and gradients were established by incubating the plates at 30°C for 24h. Supernatants were prepared as described above. *P. aeruginosa* strains were grown to early stationary phase, pelleted, and resuspended in 100 ml of MOPS buffer (10 mM MOPS, pH 7.6, and 8 mM MgSO_4_) at an OD_600_ of 1.2. 2 µl of this cell suspension was spotted approximately 6 mm from the center of the supernatant spot. Plates were incubated for 24h at 37°C followed by 24h at room temperature before images were taken of the twitching zones.

### ß-Galactosidase activity assays

To measure ß-galactosidase activity of the cAMP responsive promoter P1, 200 µl of mid-log phase *P. aeruginosa* harboring either the plasmid containing *lac*P1-*lacZ* or the empty vector were spread onto the surface of M8 + 1.5% agar, either in monoculture or coculture with equal numbers mid-log phase *S. aureus*. Cell were incubated for the indicated time points, scraped from the plates and the ß-galactosidase activity was measured as previously described (J. H. Miller 1972) and results were presented as Miller units, except that the reactions were performed at room temperature and the absorbance was measured in 96-well flat-bottom plates using a SpectraMax M2 microplate reader (Molecular Devices). For coculture conditions, in order to account for the contribution of *S. aureus* cells to the OD_600_ in the calculation for Miller Units, the OD_600_ measured for *P. aeruginosa* in monoculture was utilized. *P. aeruginosa* growth was not affected by the presence of *S. aureus*, which was confirmed by selectively plating for CFU on *Pseudomonas* Isolation Agar.

## Acknowledgements

This work was supported by funding from NIH Grant R37 AI83256 (GAO), CFF Postdoctoral Fellowship LIMOLI15F0, and CFF Postdoc-to-Faculty Transition Award LIMOLI18F5. We thank Dr. Carey Nadell for strain PA14 *attB*:: P*A1/04/03* -mKO and consultation on imaging studies, Drs. Ethan Garner, Gerard Wong, and Jeffrey Meisner for consultation on imaging studies, and Dr. Timothy Yahr for careful reading of the manuscript.

## Supplemental Figures

**Figure S1.**
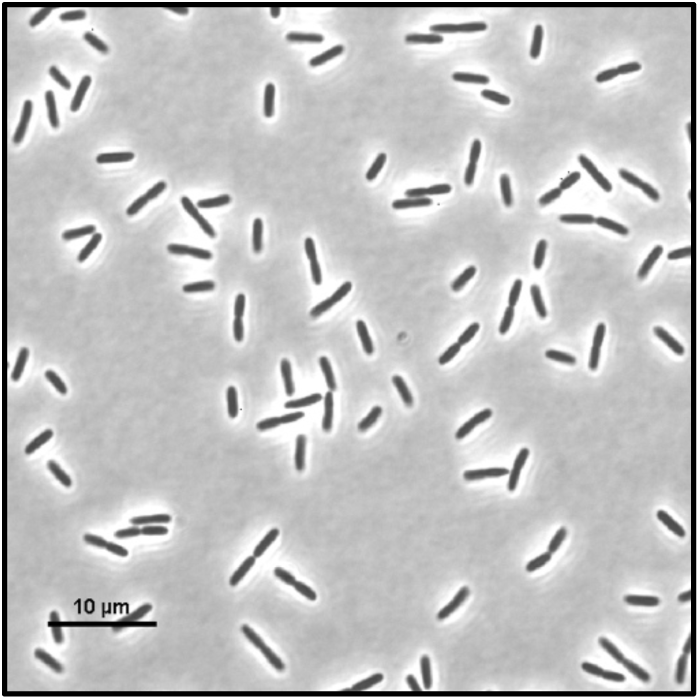
Live imaging of *P. aeruginosa* Δ*flgK* Δ*pilA* in monoculture. Representative snapshot at t=4h.

**Figure S2.**
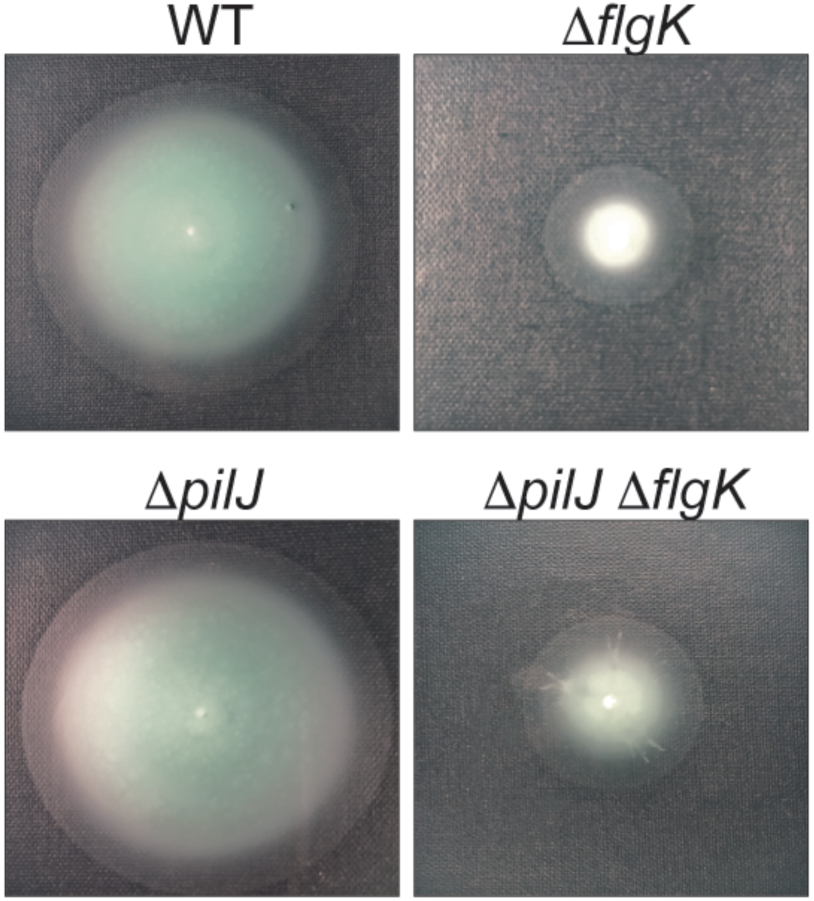
*P. aeruginosa* monoculture soft agar assay. The indicated *P. aeruginosa* strains were inoculated into soft agar (0.3%) and incubated for 24h. Representative images are shown of three independent experiments.

**Figure S3.**
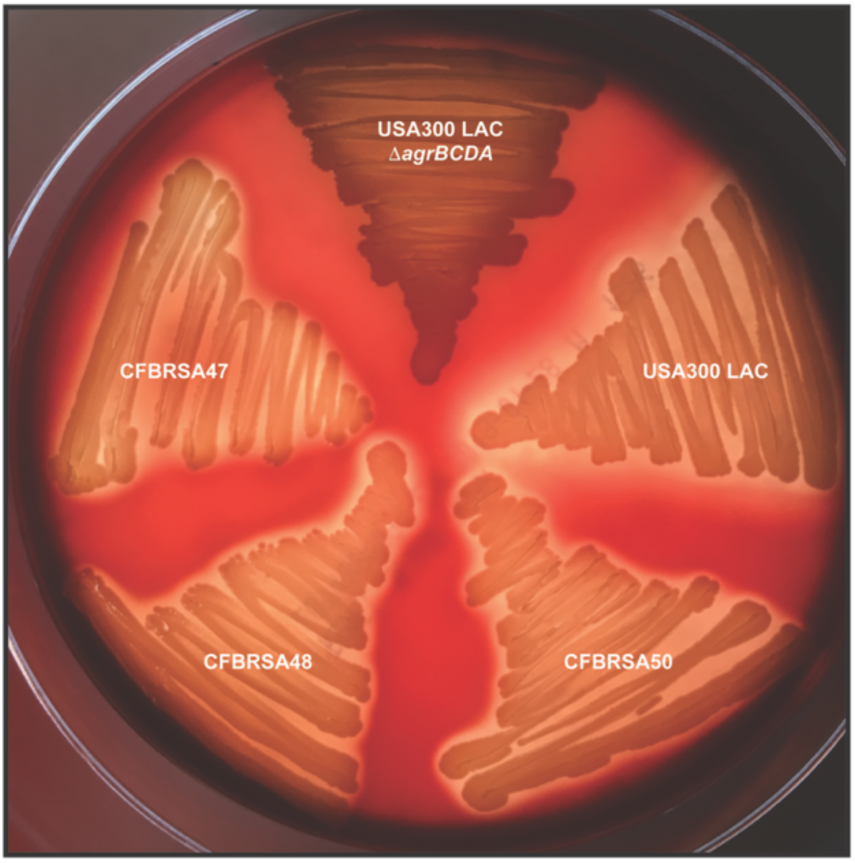
Clinical CF *S. aureus* isolates retain α-hemolysis. Laboratory *S. aureus* isolates USA300 LAC (WT) and Δ*agrBCDA* and clinical CF *S. aureus* isolates (CFBRSA47, 48, and 50) were grown on sheep’s blood agar and examined for hemolysis of red blood cells surrounding the colonies.

**Table S1.** Bacterial strains and plasmids used in this study.

| Strain or plasmid | Genotype or description | Reference or source |
| --- | --- | --- |
| *Escherichia coli* |  |  |
| DH5α | *supE44* Δ*lacU169*(ϕ80d*lacZ*ΔM15) *hsdR17 thi-1 relA1 recA1* | Life Technologies |
| S17 λpir | TpR SmR *recA*, *thi*, *pro*, *hsdR*-M+RP4: 2-Tc:Mu: Km Tn7 λpir | Life Technologies |
| *P. aeruginosa* |  |  |
| PA14 | Non-mucoid prototroph | (Rahme et al. 1995) |
| PADHL209 | PA14 *attB*:: P<sub>A1/04/03</sub>-mKO | (Nadell et al. 2017) |
| SMC3782 | PA14 Δ*pilA* | (Kuchma et al. 2010) |
| SMC5845 | PA14 Δ*flgK* | (O'Toole and Kolter 1998) |
| SMC6595 | PA14 Δ*pilA* Δ*flgK* | (Ribbe et al. 2017) |
| SMC2992 | PA14 Δ*pilJ* | (Belete et al. 2008) |
| PADHL266 | PA14 Δ*pilJ* Δ*flgK* | This study |
| SMC6852 | PA14 Δ*cyaB* | (Luo et al. 2015) |
| SMC3538 | PA14 pR::lacZ | (Wolfgang et al. 2003) |
| SMC3537 | PA14 pR::lacP1-lacZ | (Wolfgang et al. 2003) |
| CFBRPA37 | Mucoid cystic fibrosis isolate | (Limoli et al. 2017) |
| CFBRPA38 | Mucoid cystic fibrosis isolate | (Limoli et al. 2017) |
| CFBRPA43 | Mucoid cystic fibrosis isolate, *mucA22* | (Limoli et al. 2017) |
| CFBRPA44 | Non-mucoid CF isolate | (Limoli et al. 2017) |
| *S. aureus* |  |  |
| JE2 (WT) | USA300 CA-Methicillin resistant strain LAC without plasmids | (McClure and Schiller 1996) |
| SADHL07 | JE2 pCM29 (*sarAP1*-sGFP) | (Pang et al. 2010) |
| CFBRSA47 | Cystic fibrosis isolate | (Limoli et al. 2017) |
| CFBRSA48 | Cystic fibrosis isolate | (Limoli et al. 2017) |
| CFBRSA50 | Cystic fibrosis isolate | (Limoli et al. 2017) |
| SADHL44 | JE2 *agrB*::TnMar | (Fey et al. 2013) |
| SADHL82 | JE2 *sigB*::TnMar | (Fey et al. 2013) |
| SADHL87 | JE2 Δ*agrBDCA* | This study |
| SADHL91 | JE2 Δ*agrBDCA* + WT *agrBDCA* | This study |
| Plasmids |  |  |
| pMAD | *E. coli*-*S. aureus* shuttle vector. OripE194<sup>TS</sup> *ermC blaC bgaB* | (Arnaud et al. 2004) |
| pMQ30 | Shuttle vector for yeast cloning and Gram-negative allelic replacement; Gm | (Shanks et al. 2006) |

**Table S2.**
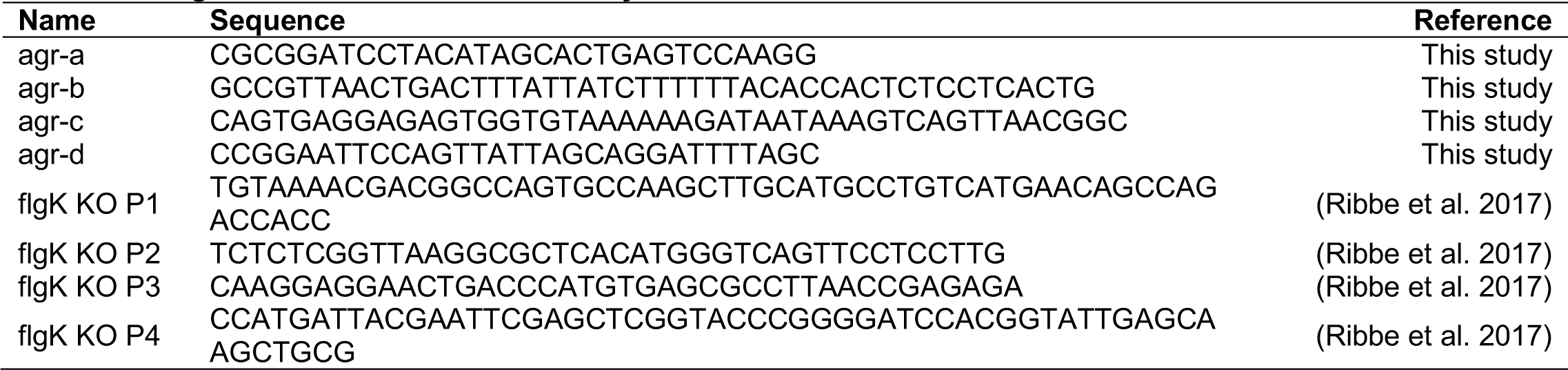
Oligonucleotides used in this study.

## Movie Legends

**Movie 1.** WT *P. aeruginosa* in monoculture. Duration 8h. Acquisition interval 15m. Playback speed 3054x.

**Movie 2.** WT *P. aeruginosa* in coculture with WT *S. aureus*. Duration 8h. Acquisition interval 15m. Playback speed 3054x.

**Movie 3.** WT *S. aureus* in monoculture. Duration 8h. Acquisition interval 15m. Playback speed 3054x.

**Movie 4.** WT *P. aeruginosa* in coculture with WT *S. aureus*. Duration 10m. 4h post inoculation. Acquisition interval 5s. Playback speed 50x.

**Movie 5.** WT *P. aeruginosa* in coculture with WT *S. aureus*. Duration 10s. 4.5h post inoculation. Acquisition interval 50 ms. Playback speed 3x.

**Movie 6.** *P. aeruginosa* Δ*pilA* in coculture with WT *S. aureus*. Duration 10s. 4.5h post inoculation. Acquisition interval 50 ms. Playback speed 3x.

**Movie 7.** *P. aeruginosa* Δ*flgK* in coculture with WT S. aureus. Duration 10s. 4.5h post inoculation. Acquisition interval 50 ms. Playback speed 3x.

**Movie 8.** *P. aeruginosa* Δ*pilA* Δ*flgK* in coculture with WT *S. aureus*. Duration 10s. 4.5h post inoculation. Acquisition interval 50 ms. Playback speed 3x.

**Movie 9.** WT *P. aeruginosa* in coculture with *S. aureus* Δ*agrBCDA*. Duration 10m. 4h post inoculation. Acquisition interval 5s. Playback speed 50x.

**Movie 10.** *P. aeruginosa* Δ*pilJ* in monoculture. Duration 10m. 4h post inoculation. Acquisition interval 5s. Playback speed 50x.

